# Zebrafish Slit2 and Slit3 act together to regulate retinal axon crossing at the midline

**DOI:** 10.1101/2022.08.12.503757

**Authors:** Camila Davison, Gabriela Bedó, Flavio R. Zolessi

## Abstract

Slit-Robo signaling regulates midline crossing of commissural axons in different systems. In the zebrafish, all retinofugal axons cross at the optic chiasm to innervate the contralateral tectum. Here, the mutant for the Robo2 receptor presents severe axon guidance defects, which were not completely reproduced in a Slit2 ligand null mutant. Since *slit3* is also expressed around this area at the stage of axon crossing, we decided to analyze the possibility that it collaborates with Slit2 in this process. We found that the disruption of *slit3* expression by sgRNA-Cas9 injection caused similar, albeit slightly milder, defects than those of the *slit2* mutant, while the same treatment in the *slit2*-/- background caused much more severe defects, comparable to those observed in *robo2* mutants. Tracking analysis of *in vivo* time-lapse experiments indicated differential but complementary functions of these secreted factors in the correction of axon turn errors around the optic chiasm. Interestingly, RT-qPCR analysis showed a mild increase in *slit2* expression in *slit3* deficient embryos, but not the opposite. Our observations support the previously proposed “repulsive channel” model for Slit-Robo action at the optic chiasm, with both Slits acting in different manners, most probably relating to their different spatial expression patterns.

## Introduction

Retinal ganglion cells (RGCs) are the projection neurons of the retina, extending their axons from the eye to the dorsal midbrain. During RGC differentiation, axons exit the eye at the optic nerve head and form the optic nerve, which transverses the ventral diencephalon toward the midline. In the zebrafish, the two nerves cross over each other to form the optic chiasm. After crossing, the optic axons form the optic tracts and continue through the diencephalon until they reach their targets in the optic tectum (Rasband et al., 2003). In certain species with binocular vision, a subset of axons does not cross at the chiasm but instead turns ipsilaterally, as is the case, for example, for mammals. When comparing different mammalian species, a correlation is found between the fraction of ipsilateral RGC projections and the degree of binocular vision: the mouse has few, while primates have many (Guillery et al., 1995). In the zebrafish, on the other hand, all RGCs project contralaterally (Rasband et al., 2003).

Axon crossing at the optic chiasm is a finely regulated process, which relies on the interplay of a wide range of signaling molecules, including Slits, Ephrins and Semaphorins. Slit molecules constitute a family of secreted glycoproteins of ≈200 kDa, with various forms present across animals with bilateral symmetry (Brose and Tessier-Lavigne, 2000). In the zebrafish, four genes have been identified (Slit1a, 1b, 2 and 3) (Hutson et al., 2003; Yeo et al., 2001). Although Slit ligands are secreted, their diffusion is limited due to their strong association with extracellular matrix components (Brose et al., 1999; Wright et al., 2012; Xiao et al., 2011). These ligands act through the Robo family of proteins, which are encoded by four genes in vertebrates, including the zebrafish (Robo1-4; (Challa et al., 2001; Lee et al., 2001; Park et al., 2003)). In addition, increasing evidence suggests that Slits can bind to other molecules, receptors or co-receptors, such as heparan sulfate proteoglycan (HSPG) (Steigemann et al., 2004) or PlexinA1, a Semaphorin receptor (Delloye-Bourgeois et al., 2015).

The Slit-Robo signaling pathway has been proven instrumental for commissural axon crossing at the midline (Herrera et al., 2019). Slit2 is the most studied in this regard, and was shown to be essential in guiding axons in a wide variety of neuronal types, including RGCs (Niclou et al., 2000; Ringstedt et al., 2000). In the zebrafish, we previously showed that loss of *slit2* causes defects in axon organization at the optic nerve, optic chiasm and the proximal portion of the optic tract, as well as minor guidance defects and a reduction in growth cone velocity around the midline (Davison and Zolessi, 2021). However, this can be considered a relatively “mild” phenotype when compared to that observed in *astray* embryos, mutant for the putative receptor expressed in RGCs, Robo2 (Fricke et al., 2001). This suggests the presence of other Slits regulating the midline crossing of retinal axons. A strong candidate for this is Slit3, whose mRNA is expressed around the optic chiasm at the same time as that of Slit2, albeit with a different spatial distribution (Chalasani et al., 2007). Even though the role of Slit3, acting either by itself or together with Slit2, has been previously analyzed in zebrafish commissural axons, such as those of the supraoptic tract (SOT; (Zhang et al., 2020)) or the postoptic commissure (POC; (Barresi et al., 2005)), the functional interaction of these two ligands at the optic chiasm remains unexplored.

In this work, we analyzed the hypothesis that both Slit2 and Slit3 might cooperate to organize retinal axons at the optic chiasm by generating zebrafish embryos deficient for these factors using CRISPR/Cas9 technology. Interestingly, we were able to partially reproduce the phenotype of Robo2-deficient embryos when disrupting *slit2* and *slit3* simultaneously. This was evidenced as the appearance of severe axon guidance errors around the chiasm, including invasion of extra-optic regions or misdirections in the optic pathway, and innervation of the ipsilateral optic tectum with the formation of a secondary commissure. Time-lapse imaging of growing axons also revealed some apparent functional differences of *slit2* and *slit3* in regulating RGC axon crossing at the midline. The milder defects seen in the single-deficient embryos could be partly due to these functional differences, probably based on their different expression patterns, as mentioned above (Chalasani et al., 2007; Davison and Zolessi, 2021), and partly to a modest but significant compensation of *slit3* deficiency by an increase in *slit2* expression. Altogether, our observations support the “repulsive channel” model for the action of Slits at the brain commissures (Hutson and Chien, 2002), and suggest that both Slit2 and Slit3 could be acting through the Robo2 receptor to guide retinal axons at the optic chiasm.

## Materials and methods

### Fish breeding and care

Zebrafish were maintained and bred in a stand-alone system (Tecniplast), with controlled temperature (28 °C), conductivity (500 μS/cm^2^) and pH (7.5), under live and pellet dietary regime. Embryos were raised at temperatures ranging from 28.5 to 32 °C and staged in hours post-fertilization (hpf) according to Kimmel and collaborators (Kimmel et al., 1995). Fish lines used: wild-type (Tab5), Tg(*atoh7*:gap43-EGFP)^cu1^ (Zolessi et al., 2006), *robo2* mutant *astray*^*ti272z*^ (Karlstrom et al., 1996) and the CRISPR-generated mutant line NM_131753.1:g.30_39del, or *slit2*-/-^*ipm1*^ (Davison and Zolessi, 2021). Since the homozygotes are viable in this mutant line, for the present report we have used only maternal-zygotic mutants, hereafter referred to as *slit2-/-*^*mz*^. All the manipulations were carried out following the approved local regulations (CEUA-Institut Pasteur de Montevideo, and CNEA).

### Embryo microinjection

We designed four single-guide RNAs (sgRNA) against the *slit3* gene using the CRISPRscan tool (Moreno-Mateos et al., 2015) and injected them together with mRNA for the zfCas9 flanked by two nuclear localization signal sequences (“nCas9n”), previously reported as highly efficient (Jao et al., 2013). We first tried each sgRNA individually to assess their toxicity and efficiency. We recognized one sgRNA complementary to a sequence in the second exon of the *slit3* gene (*slit3* 206, indicating the position of its binding site in the coding sequence; Table S1) as having no toxic effects and being highly efficient based both on microscopic inspection of the phenotypes and genotyping of the expected mutation site in 72 hpf embryos (as described below). We injected one-cell stage Tab5 or *slit2-/-*^*mz*^ embryos with this sgRNA, together with nCas9n mRNA. Alternatively, we co-injected the *slit3* sgRNA together with a *slit2* sgRNA previously reported by us (*slit2* 71; Table S1 (Davison and Zolessi, 2021)). For mosaic transgenic labeling, 2.5 pg of DNA coding for *atoh7*:EGFP-CAAX (Lepanto, 2017) were injected, together with 6 pg of Tol2 transposase mRNA.

### Embryo genotyping

For genotyping, genomic DNA from single embryos was extracted at 72 hpf. Individual embryos were placed in tubes and 20 μL of 50 mM NaOH was added. After a 15 min incubation at 95° C, the tubes were put on ice and briefly homogenized using a P-20 micropipette. Finally, 2 μL of 1 M Tris-HCl (pH 8) were added, followed by a 1 min centrifugation at 20000 g. The resulting supernatant was used for the PCR reaction (2.5 μL per 10 μL reaction), with the specific primers listed in Table S2, flanking the target sequence of the *slit3* 206 sgRNA. PCR conditions were as follows: initial denaturation step at 95 °C for 2 min, followed by 30 cycles at 95 °C for 30 s, 60 °C for 30 s and 72 °C for 2 min, and a final extension at 72° C for 7 min. The PCR products obtained were analyzed through electrophoresis on either 3% agarose or 8% polyacrylamide gels.

### RNA isolation and real-time quantitative RT-PCR

Total RNA was extracted from the cephalic region of 30 and 48 hpf embryos. For each stage, four conditions were analyzed: non-injected wild-type, *slit2-/-*^*mz*^, *slit3* crispants, and *slit2-/-*^*mz*^*;slit3* crispant embryos. 40 embryos (coming from at least three different crosses) corresponding to each condition were dissected, lysed and the heads were stored in Trizol at -80 °C. Total RNA was isolated using the Direct-zol RNA MiniPrep kit (Zymo Research), following the manufacturer’s protocol. cDNA was synthesized from normalized RNA amounts using the RevertAid First Strand cDNA Synthesis kit (Thermo Fisher), with a mixture of oligo dT and random primers. Quantitative PCR was performed using the SensiFAST Sybr Low Rox kit (Bioline) with specific primers listed in Table S2. The qPCR was done in a 7500 Real-Time PCR System (Applied Biosystems), and the conditions were as follows: initial denaturation step at 95 °C for 2 min, followed by 40 cycles at 95 °C for 10 s, 60 °C for 15 s and 72 °C for 30 s.

Two biological samples and two technical replicas were used. Expression fold-changes between groups were calculated relative to internal control expression (*eIF1a*), according to Pfaffl et al. (Pfaffl et al., 2002). Statistical significance was determined using a permutation-based procedure, the non-parametric pairwise fixed reallocation randomization test, implemented in the Relative Expression Software Tool (REST). P value < 0.05 was considered statistically significant.

### Whole-mount immunofluorescence

Embryos were grown in 0.003% phenylthiourea (Sigma, St. Louis, USA) from 10 hpf onwards to delay pigmentation, and fixed overnight at 4 °C, by immersion in 4% paraformaldehyde in phosphate buffer saline, pH 7.4 (PFA-PBS). For whole-mount immunostaining all subsequent washes were performed in PBS containing 1% Triton X-100. Further permeability was achieved by incubating the embryos in 0.25% trypsin-EDTA for 10–15 min at 0 °C. Blocking was for 30 min in 0.1% bovine serum albumin (BSA), 1% Triton X-100 in PBS. The primary antibody zn8 (ZIRC, Oregon, USA), recognizing the adhesion molecule neurolin/DM-grasp, was used 1/100 in blocking solution. The secondary antibody used was anti-mouse IgG-Alexa 488 (Life Technologies, Carlsbad, USA), 1/1000 in blocking solution. Nuclei were fluorescently stained with methyl green (Prieto et al., 2014). All antibody incubations were performed overnight at 4 °C. Embryos were mounted in 1.5% agarose-50% glycerol in 20 mM Tris buffer (pH 8.0) and stored at 4 °C or −20 °C. Observation of whole embryos was performed using a Zeiss LSM 880 laser confocal microscope, with a 25x 0.8 NA glycerol immersion objective.

### Lipophilic dye labeling

Phenylthiourea-treated embryos were fixed at 48 hpf or 5 dpf as described above. They were then immobilized on glass slides using 2% agarose and injected with 1,1’-dioctadecyl-3,3,3′,3′-tetramethylindocarbocyanine perchlorate (DiI, Molecular Probes) or 3,3′-dioctadecyloxacarbocyanine perchlorate (DiO, Molecular Probes) dissolved in chloroform. For optic chiasm observation in 48 hpf embryos, DiI was injected into the vitreous chamber of one eye, whereas DiO was injected into the contralateral eye. For optic tectum observation in 5 dpf larvae, DiI was injected into the vitreous chamber of one eye. In all of the cases, after the injection the embryos or larvae were incubated for 48 hours at room temperature and the dissected brains were mounted in 1.5% agarose-50% glycerol in 20 mM Tris buffer (pH 8.0) and stored at 4 °C or -20 °C. Observation was performed using a Zeiss LSM 880 laser confocal microscope, with a 25× 0.8 NA glycerol immersion objective.

### Time-lapse imaging

Embryos were selected around 30 hpf, anesthetized using 0.04 mg/mL MS222 (Sigma) and mounted in 1% low-melting point agarose (Sigma) over glass bottom dishes. After agarose gelification and during image acquisition, embryos were kept in Ringer’s solution (116 mM NaCl, 2.9 mM KCl, 1.8 mM CaCl2, 5 mM HEPES, pH 7.2) with 0.04 mg/mL MS222. Live acquisitions were made using a Zeiss LSM 880 laser confocal microscope, with a 40X, 1.2 NA silicone oil immersion objective. Stacks around 40 μm-thick were acquired in bidirectional scanning mode, at 1 μm spacing and 512×512 pixel resolution every 15 minutes, for 2.5-16.5 hours. The acquisition time per embryo was approximately 1 min, and up to 8 embryos were imaged in each experiment. The embryos were fixed in 4% PFA immediately after the end of the time-lapse, and processed for further confocal microscopy, labeling nuclei with methyl green.

### Microscopy data analysis

Images were analyzed using Fiji (Schindelin et al., 2012). Tracking of axons to obtain distance and velocity measurements was performed with the Manual Tracking plugin (https://imagej.net/Manual_Tracking), using the site of pioneer axon crossing at the midline as a reference point. For all these analyses, the embryo midline was set as 0 in “x” (the medio-lateral dimension) and the ventral surface of the diencephalon as 0 in “y” (the ventro-dorsal dimension). The instantaneous velocity of the growth cone was determined for each point and expressed in μm/min, and the turn angle (difference between consecutive axon growth direction angles with respect to the “x” axis) was calculated and expressed in degrees (°). In order to better compare between these angles, we applied a correction for the curvature based on the averaged angles in the control embryos (subtracting 0.5° to each value). To visualize and quantify the eventual “axon turn errors”, we used the absolute values of the turn angles, and only analyzed those values that were larger than the mean value in control embryos (10.4°).

Statistical analysis was performed using GraphPad Prism software. As a routine, the data sets were checked for normality using Kolmogorov-Smirnoff normality test. For multiple sets of data, in the case of normal distributions we used either uncorrected one-way ANOVA or Brown-Forsythe and Welch test, for equal or unequal SD values respectively, or the Kruskal-Wallis test for non-normal distributions. For the comparison of proportions, we used the Fisher’s exact test.

## Results

### Simultaneous loss of Slit2 and Slit3 causes retinal axon misprojections at the optic chiasm comparable to those observed upon Robo2 loss

To determine if *slit3* has a role in retinal axon guidance at the midline, we set out to generate F0 mosaic mutants using the CRISPR/Cas9 system (crispants). We identified an sgRNA (*slit3* sgRNA 206; Table S1) complementary to a sequence in the second exon of the *slit3* gene which caused indels in the target genomic sequence with high efficiency, as evidenced by the appearance of diffuse or extra bands in the electrophoresis analysis of PCR products (Fig. S1). The injection of *slit3* sgRNA/nCas9n in wild-type embryos resulted in a phenotype characterized by the appearance of some cases of bifurcation of one optic nerve around the other at the optic chiasm, as evidenced through zn8 immunostaining of RGCs (Fig. 1A and Video S1). A similar phenotype was observed in non-injected *slit2-/-*^*mz*^ mutantembryos (Fig. 1A and Video S1), as previously described (Davison and Zolessi, 2021). In both *slit3* crispants and in *slit2-/-*^*mz*^ mutants, this phenotype appeared with similar a penetrance of around 25% (Fig. 1B). Moreover, we found that the two branches resulting from nerve bifurcation could be symmetrical or asymmetrical, depending on the proportion of axons going into each branch, with an apparently equal occurrence of thinner branches on the anterior or posterior side (Fig. S2). There were no significant differences in the proportion of symmetrical/asymmetrical bifurcations between *slit2-/-*^*mz*^ and *slit3* crispants (Fig. S2). Surprisingly, when we injected *slit3* sgRNA/nCas9n in *slit2-/-*^*mz*^ embryos, we observed a relatively high frequency of new, more severe, defects at the optic chiasm, together with a marked reduction in the penetrance of the nerve bifurcation phenotype to around 10% (Fig. 1A-C). These new defects largely consisted of anterior misprojections of retinal axons at the optic chiasm, similar to what is observed in *astray/robo2* mutant embryos, and that were never detected in *slit2-/-*^*mz*^ mutant or *slit3* crispant embryos (Fig. 1A,C,D and Video S1).

**Figure 1.**
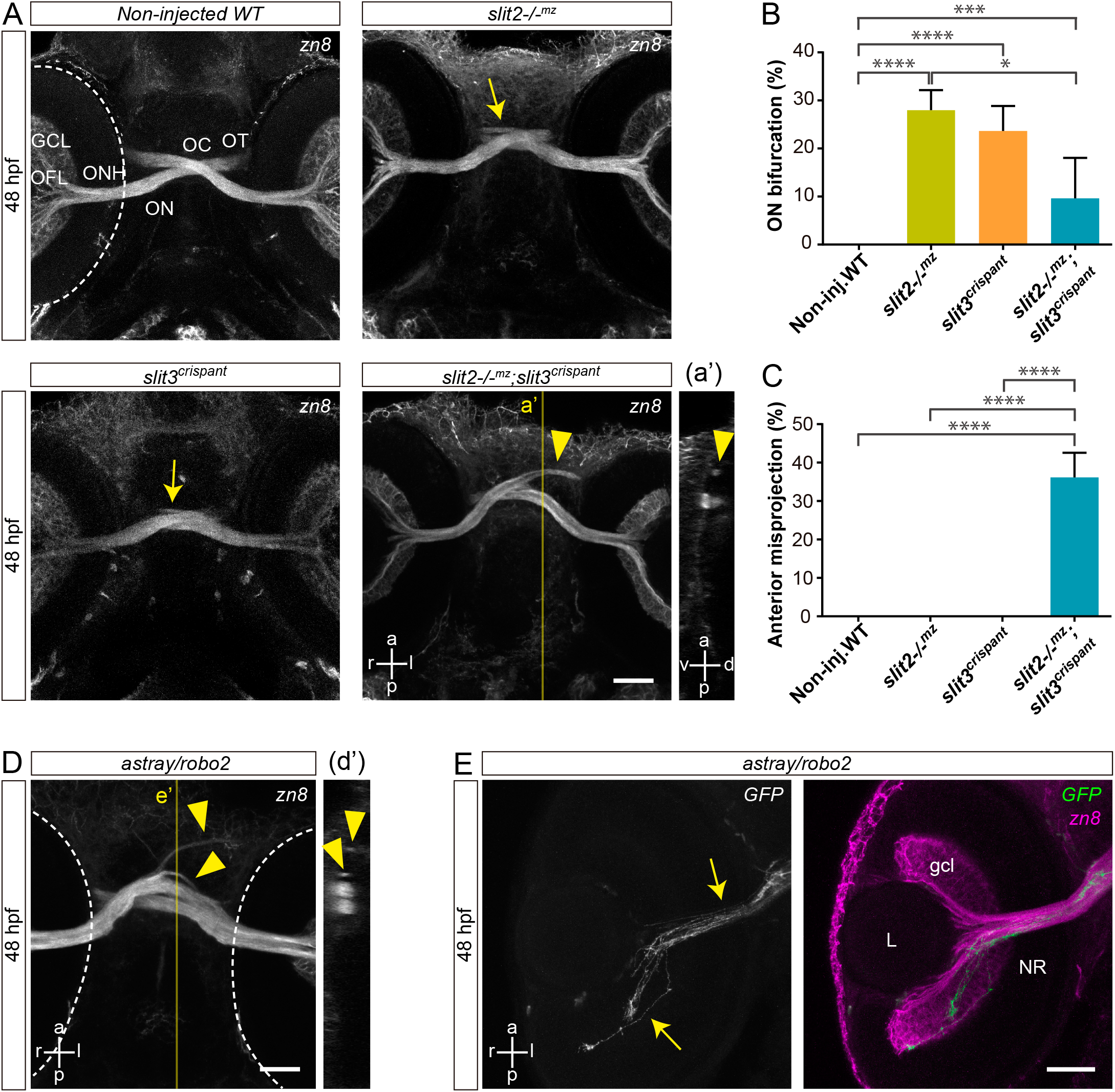
Simultaneous loss of Slit2 and Slit3 causes retinal axon misprojections at the optic chiasm I: gross phenotype and comparison with Robo2 loss. **A**. Maximum intensity z-projections of the cephalic region of 48 hpf embryos immunostained to label RGCs (zn8 antibody; ventral view). Optic nerve bifurcation can be observed in *slit2-/-*^*mz*^ embryos and *slit3* crispants (arrows). Injection of *slit3* sgRNA/nCas9n in *slit2-/-*^*mz*^ embryos results in more severe defects at the optic chiasm, including anterior projections of the optic nerves (arrowhead), clearly observed in the orthogonal section in a’. **B**,**C**. Penetrance of the bifurcation phenotype (B) and the anterior projection phenotype (C). n embryos (n independent experiments) = 81 (3) non-injected controls, 53 (3) *slit2-/-*^*mz*^, 57 (3) *slit3* crispants, 70 (3) *slit2-/-*^*mz*^;*slit3* crispants; mean ± SD of the 3 experiments is shown; level of statistical significance shown as asterisks, based on a p < 0.05 and analyzed by Fisher’s exact test for proportions. **D**. Maximum intensity z-projection of the cephalic region of a 48 hpf *astray/robo2* embryo immunostained to label RGCs (zn8 antibody; ventral view). Defects can be observed at the optic chiasm, including anterior projections of the optic nerves (arrowhead), clearly evidenced in the orthogonal section in d’. **E**. Maximum intensity z-projection of the retina of a 48 hpf *astray/robo2* embryo injected with *atoh7*:EGFP-CAAX DNA, where only the contralateral retina was labeled. Axons from labeled RGCs can be seen growing into the contralateral eye through the optic nerve (arrows). Dashed lines: outer retinal borders. GCL: ganglion cell layer; L: lens; NR: neural retina; OC: optic chiasm; OFL: optic fiber layer; ON: optic nerve; ONH: optic nerve head; OT: optic tract. Scale bars: A, 40 μm; D, E, 30 μm. See Video S1.

*astray/robo2* mutants also presented misguided RGC axons growing into the contralateral optic nerve, and even reaching the contralateral retina, as evidenced in mosaically-labeled embryos displaying unilateral *atoh7*:EGFP-CAAX expression (Fig. 1E). To better characterize the axon navigation errors observed upon *slit2* and/or *slit3* expression disruption, we differentially labeled the optic nerves using the lipophilic dyes DiI and DiO. As described for zn8 labeling, we found clear evidence of a proportion of embryos with a bifurcated optic nerve in *slit2-/-*^*mz*^, clearly contrasting with the complete segregation of the two optic nerves at the chiasm in wild-type embryos (Fig. 2A,B and Video S2). This phenotype was also present in a similar proportion of *slit3* crispants (Fig. 2C and Video S2). On the other hand, *slit2-/-*^*mz*^*;slit3* crispants showed a severe alteration of axon organization at the chiasm. In some cases, one of the optic nerves split around the contralateral nerve, but the branches were not observed to re-connect at the optic tract (Fig. 2E and Video S2). Furthermore, misprojections of retinal axons were observed, both into the ipsilateral optic tract and into the contralateral optic nerve (Fig. 2D,E and Video S2). These observations are summarized in Fig. 2F.

**Figure 2.**
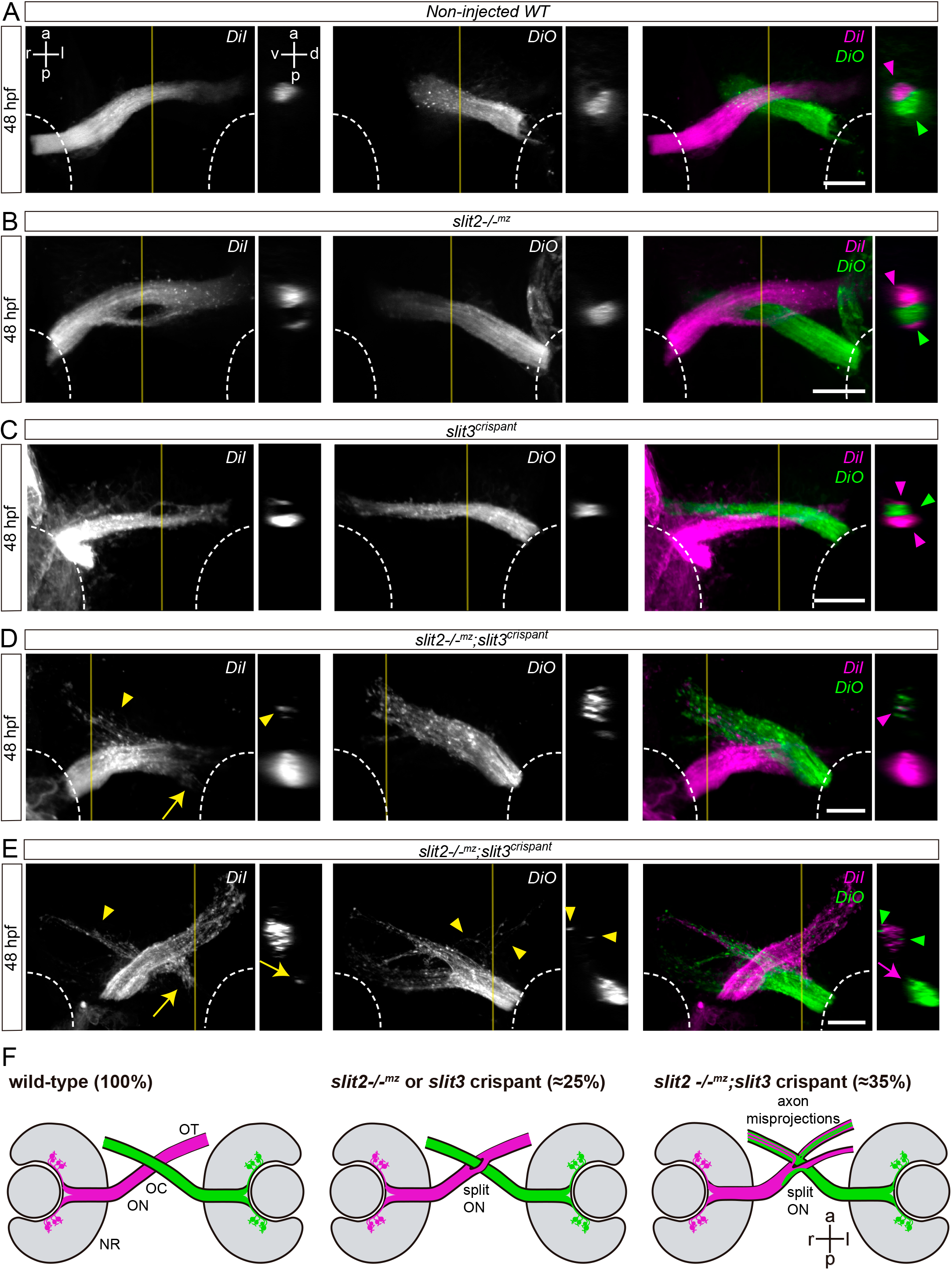
Simultaneous loss of Slit2 and Slit3 causes retinal axon misprojections at the optic chiasm II: anterograde axon labeling analysis. RGC axons from both eyes were anterogradely labeled through DiI or DiO injection into the retina, and images show the optic chiasm region (ventral view). **A-E**. Maximum intensity z-projections of the midline region of 48 hpf of non-injected wild-type (A), *slit2-/-*^*mz*^ (B), *slit3* crispant (C) and *slit2-/-* ^*mz*^;*slit3* crispant (D,E) embryos. Arrowheads: projections into the ipsilateral optic tract. Arrows: projections into the contralateral optic nerve. **F**. Diagrams summarizing the main phenotypic features of each experimental situation shown in this and the previous figures. NR: neural retina; OC: optic chiasm; ON: optic nerve; OT: optic tract. Scale bars: 30 μm. See Video S2.

### RGCs in slit2/slit3 double crispants partially project to the ipsilateral optic tectum

To assess whether the ipsilateral retinal projections we observed at the optic chiasm of *slit2, slit3* and *slit2+slit3* deficient embryos reach the optic tectum, we decided to follow axon trajectories using lipophilic dye tracing. For this, we injected DiI into the right eye of fixed whole-mount 5 dpf *atoh7*:Gap-EGFP transgenic larvae, in which RGC axons from both eyes are labeled with membrane-tagged EGFP. After eye removal, axonal projections to the contralateral and ipsilateral optic tecta were analyzed in dorsal view (Fig. 3). We observed no DiI-labeled axons at the ipsilateral optic tectum of wild-type, *slit2* or *slit3* crispant larvae (Fig. 3A-C). However, in *slit2+slit3* crispants, even though most DiI-positive axons innervated the contralateral tectum, a small proportion of them extended into the ipsilateral midbrain (Fig. 3D,d’ and Video S3). This phenotype was observed in 62.5% of these double-deficient larvae (n = 10/16). In all ipsilateral innervation cases there were axons ending at the pretectal arborization field 9 (AF-9), while in most (n = 8/10) there were also some axons innervating the ipsilateral optic tectum. Interestingly, some of the ipsilateral-innervating axons seemed to cross the midline at a site different from the optic chiasm, which localized slightly posterior to it (Fig. 3D).

**Figure 3.**
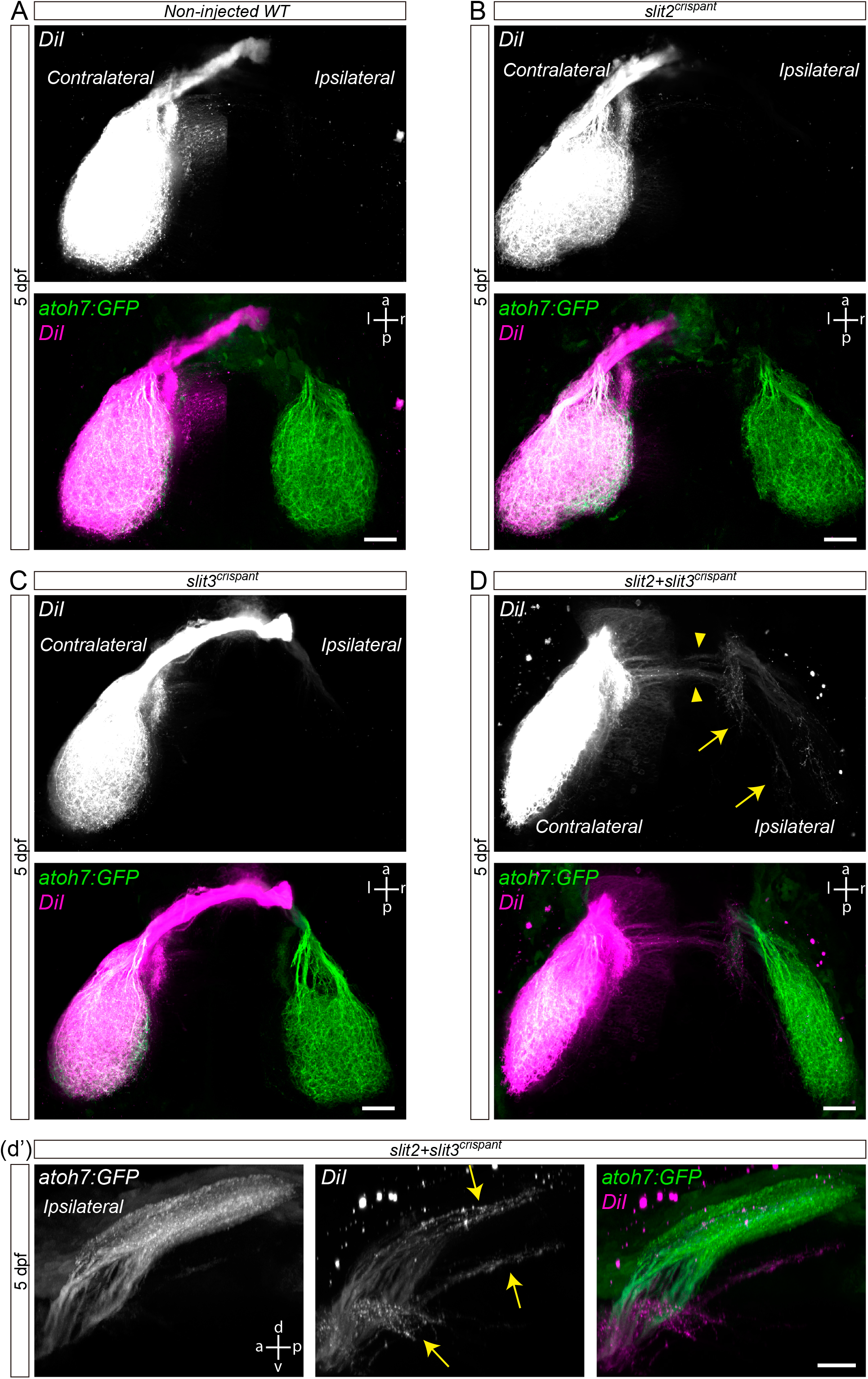
Retinal axons from *slit2+slit3* crispants innervate the ipsilateral optic tectum. Retinotopic anterograde RGC axon DiI labeling in *atoh7*:Gap43-EGFP (*atoh7*:GFP) transgenic larvae. **A-D**. Horizontal maximum intensity z-projections of the cephalic region of 5 dpf larvae, where both the contralateral and ipsilateral tecta can be observed (dorsal view). Non-injected wild-type (A), *slit2* crispant (B), *slit3* crispant (C) and *slit2+slit3* crispant (D) larvae are shown. DiI-labeled axons are observed in the ipsilateral tectum of *slit2+slit3* crispants (arrows), and can be better visualized in the rotated view in d’. DiI-labeled axons can also be observed crossing the midline at a site slightly posterior to the optic tract (arrowheads). Scale bars: A-E, 30 μm; d’, 20 μm. See Video S3.

### slit2 expression is upregulated in slit3 crispants at early stages

In order to determine whether a mutual genetic compensation between *slit2* and *slit3* might contribute to the milder effects observed in the single-deficient embryos, we performed quantitative RT-PCR analyses for *slit2* and *slit3* expression in non-injected wild-type, *slit2-/-*^*mz*^, *slit3* crispant and *slit2-/-*^*mz*^*;slit3* crispant embryos, at both 30 and 48 hpf. Fold-change in gene expression for each condition and developmental stage are shown in Table 1. Although no change in *slit3* expression was detected in *slit2* mutants, a significant up-regulation of *slit2* was evident in *slit3* crispants at 30 hpf, indicating a genetic compensation phenomenon. This increase in gene expression was no longer detected at 48 hpf, when the optic chiasm has already formed.

**Table 1.** Quantitative RT-PCR analysis of gene expression for *slit2* and *slit3* in zebrafish embryo heads. Comparison is expressed as the fold-change between the test and the wild-type conditions. Values were calculated using REST, and p < 0.05 was considered statistically significant. Significant increase is shown in bold, and decrease in italics.

| Gene | Test | 30 hpf |  | 48 hpf |  |
| --- | --- | --- | --- | --- | --- |
|  |  | Fold-change | p | Fold-change | p |
| *slit2* | *slit2*<sup>-/-mz</sup> | *0.450* | 0.001 | *0.337* | 0.001 |
|  | *slit3*<sup>crispant</sup> | **1.357** | 0.001 | 1.050 | 0.277 |
|  | *slit2*<sup>-/-mz</sup>;*slit3*<sup>crispant</sup> | *0.452* | 0.001 | *0.525* | 0.008 |
| *slit3* | *slit2*<sup>-/-mz</sup> | 0.915 | 0.394 | 1.002 | 0.950 |
|  | *slit3*<sup>crispant</sup> | 0.932 | 0.646 | *0.727* | 0.016 |
|  | *slit2*<sup>-/-mz</sup>;*slit3*<sup>crispant</sup> | *0.656* | 0.003 | *0.652* | 0.008 |

Although the indels generated by CRISPR/Cas9 using a single sgRNA directed to the protein-coding sequence are not expected to affect mRNA transcription of the mutated gene, we also analyzed the expression of each gene in their respective deficient situation. We previously showed that the mutation in *slit2-/-*^*mz*^ embryos caused a marked reduction in the amount of mRNA detected by RT-PCR (Davison and Zolessi, 2021). This effect is actually not unusual, and is most likely mediated by the cellular mechanism known as nonsense-mediated mRNA decay (Karousis and Mühlemann, 2019). Here, we again detected a significant *slit2* mRNA decrease by RT-qPCR at 30 and 48 hpf in the *slit2-/-*^*mz*^ mutants, both non-injected and injected with the *slit3* sgRNA (Table 1). In the case of *slit3* expression in *slit3* crispants, a smaller but significant decrease was also evident at 48 hpf, both in the wild-type and *slit2-/-*^*mz*^ background (Table 1). At 30 hpf, however, collected data appeared more variable among the analyzed samples, resulting in a significant reduction of *slit3* expression in *slit2-/-*^*mz*^-*slit3* crispant embryos, but not in *slit3* crispants.

### Slit2 and Slit3 deficiency increase minor axon guidance defects in RGC axons crossing the optic chiasm

With the aim of better understanding the growth dynamics of retinal axon crossing in embryos deficient for *slit2* and/or *slit3*, we decided to follow the growth of axons across the optic chiasm by time-lapse microscopy. We performed these experiments by injecting *slit2* and *slit3* sgRNA/nCas9n, either individually or together, in *atoh7*:Gap-EGFP transgenic embryos (Fig. 4 and Videos S4-8). In non-injected controls, axons usually extended quickly from the optic nerve through the chiasm, showing very little deviation from a relatively smooth curve (Fig. 4A). On the other hand, axons from *slit2, slit3* and *slit2+slit3* crispants showed more sinuous pathways, together with several axon guidance defects (Fig. 4B-D). In all three conditions, we observed ipsilateral turns and axon defasciculation, both of which occurred before and after the growth cone had crossed the midline. We quantified the proportion of embryos in which defects were observed, and we found that, even though these were present in *slit2, slit3* and *slit2+slit3* crispants, the frequency was slightly higher in *slit2+slit3* crispant embryos (Table 2). It is important to note that this quantification was performed on both pioneer and follower axons.

**Table 2.** Quantification of axon guidance defects. The analysis was performed on the time-lapse experiments used for axon growth cone tracking. Pathfinding errors were classified into four categories, and representative examples of each category are shown in Fig. 4. The frequency of these errors was quantified considering both pioneer and follower axons. Asterisks indicate a proportion significantly different to that of non-injected wild-type embryos, according to the Fisher’s exact test.

|  | Non-injected<br>WT<br>(n=13) | *slit2*<br>crispants<br>(n=14) | *slit3*<br>crispants<br>(n=16) | *slit2+slit3*<br>crispants<br>(n=12) |
| --- | --- | --- | --- | --- |
| Projections to ipsilateral tract (%) | 7.7 | 14.3 | 12.5 | 33.3 |
| Projections to contralateral nerve (%) | 0.0 | 0.0 | 0.0 | 8.3 |
| Fasciculation defects (%) | 0.0 | 42.9 * | 25.0 | 58.3 ** |
| Minor pathfinding errors (%) | 0.0 | 21.4 | 12.5 | 41.6 * |
| No defects (%) | 92.3 | 42.9 * | 62.5 | 33.3 ** |

**Figure 4.**
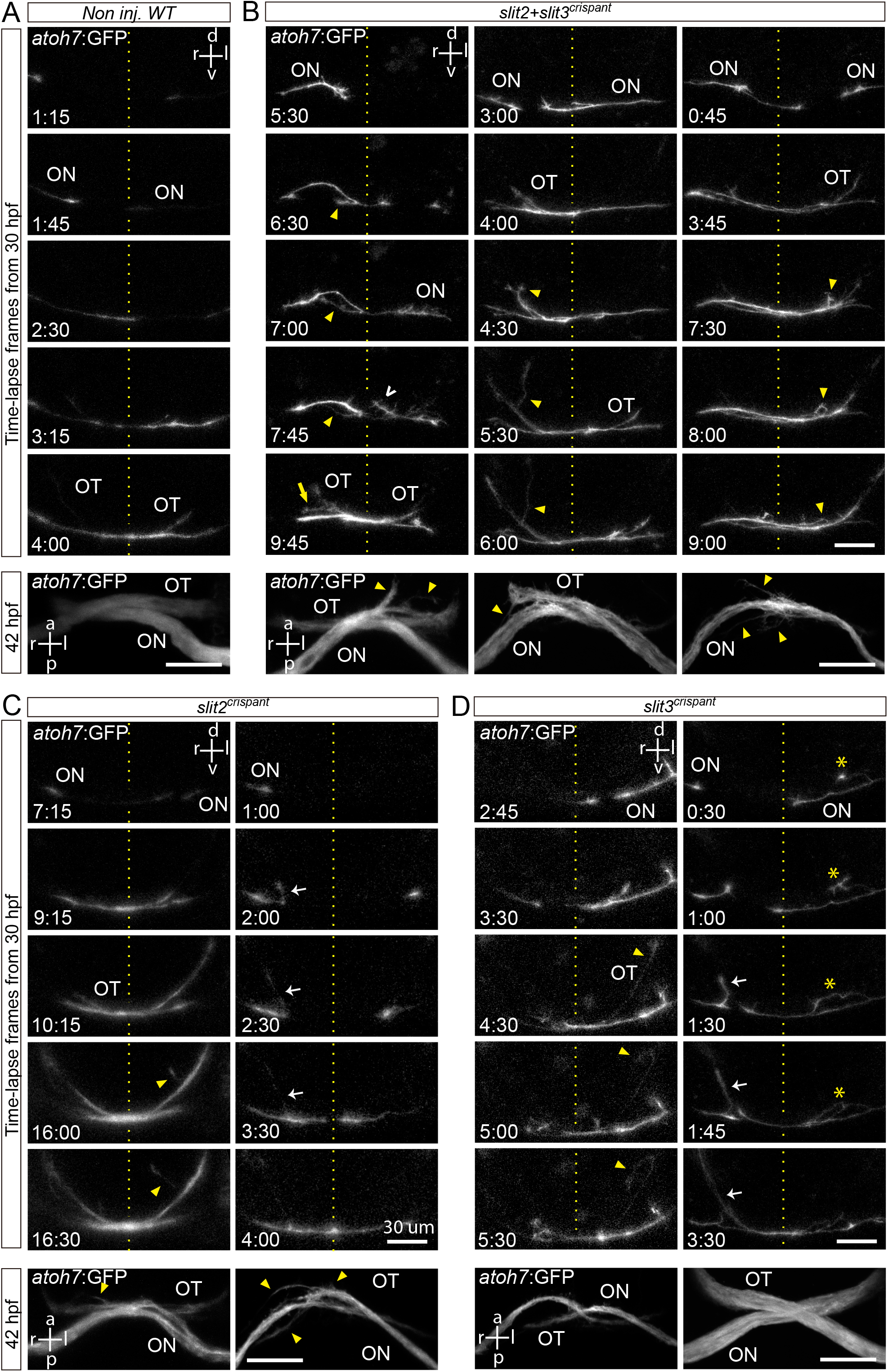
Retinal axons from *slit2, slit3* and *slit2/slit3* double crispants present navigation errors around the midline. Confocal time lapse observations of pioneer optic axon growth along the area surrounding the optic chiasm, in *atoh7*:Gap43-EGFP (*atoh7*:GFP) transgenic embryos. **A-D**. Sequences of selected frames from time-lapse analysis of non-injected wild-type (A), *slit2+slit3* crispant (B), *slit2* crispant (C) and *slit3* crispant (D) embryos. A frontal view of the chiasm area is shown, starting with the appearance of the first axon at the optic nerve. The dashed lines indicate the location of the midline. Maximum intensity z-projections of the corresponding embryos are shown at the bottom of each time-lapse sequence, in ventral view. Misrouted axons were classified and quantified (Table 2) as follows: projections to the ipsilateral optic tract (e.g., white arrows), projections to the contralateral optic nerve (e.g., yellow arrow), fasciculation defects (e.g., asterisks), and immediately corrected minor pathfinding errors (e.g., empty white arrowhead). Yellow arrowheads point to other errors seen in these images. ON: optic nerve; OT: optic tract. Scale bars: 30 μm. See Videos S4-S8.

### Axon trajectory and instantaneous velocities are differentially affected upon Slit2 and Slit3 expression disruption

We then decided to further analyze the described defects on axon growth by digitally tracking the path of only pioneer axon growth cones, which would not be affected by axon-axon fasciculation signaling (Fig. 5). When axon trajectories are compared in this way, in addition to an important variability between individual axons/embryos, some qualitative differences are evident between the different experimental situations. The most remarkable deviations from the control situation were observed in the *slit2* crispant embryos, at the level of the proximal optic tract. Here, axon trajectories appeared more disperse in general, and several axon turn errors were evident as sudden changes in direction (Fig. 5A). The *slit3* and the double crispants did not present such an evident difference to the wild-type condition, although the double crispants surprisingly showed reduced errors and dispersion in the optic tract area when compared with the *slit2* crispants (Fig. 5A).

**Figure 5.**
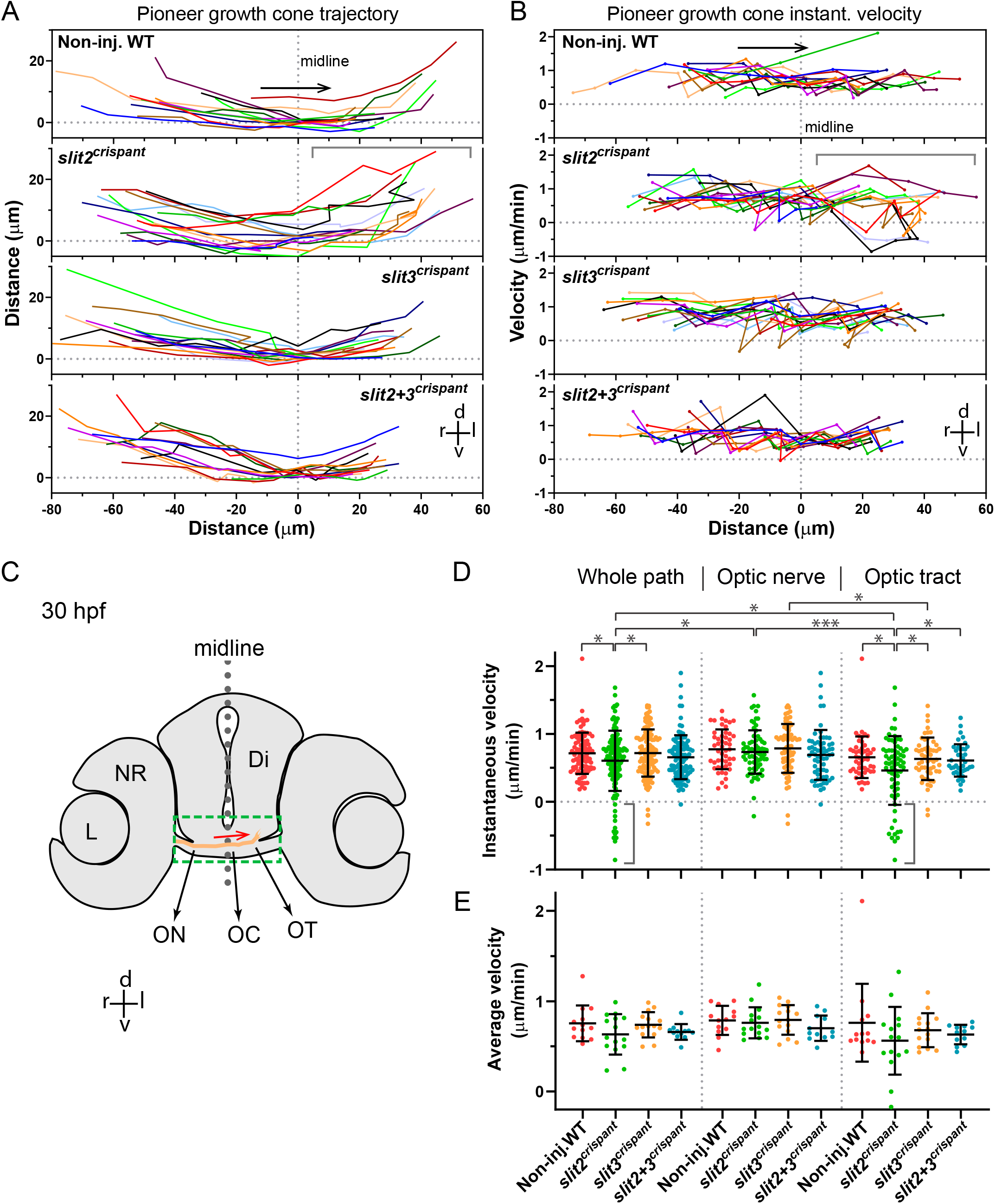
Tracking of pioneer axon growth through the optic chiasm highlights differential effects of Slit2 and Slit3 loss I: analysis of axon trajectories and velocities. 2D tracking analysis of RGC axon growth along the optic nerve, optic chiasm and proximal optic tract, from time-lapse experiments starting at 30 hpf, shown in Fig. 4. **A**. Axon trajectories across the ventro-dorsal and medio-lateral axes, for non-injected and crispant embryos. The gray bracket marks an area of higher spatial dispersion and/or apparent axon growth deviations in the *slit2* crispants. **B**. Instantaneous growth cone velocities compared to the medio-lateral axis along the optic pathway. The gray brackets mark areas of higher velocity dispersion and/or apparent axon growth deviations in the *slit2* crispants. **C**. Diagram showing a cross-section of a 30 hpf zebrafish embryo at the level of the optic chiasm, highlighting the approximate area analyzed in the time-lapse tracking experiments, with an example of the trajectory of a wild-type axon. Di: diencephalon; L: lens; NR: neural retina; OC: optic chiasm; ON: optic nerve; OT: optic tract. **D-E**. Instantaneous velocities for the four experimental situations, analyzed as a pool (D) or averaged per embryo (E). In all cases, the data were analyzed and compared either for the whole axon path tracked, the optic nerve portion or the optic tract portion, as noted at the top. The gray brackets in D highlight several negative velocity values in the *slit2* crispants. Mean ± SD; statistical significance is shown as asterisks, based on a p < 0.05 and analyzed by the Brown-Forsythe and Welch test for multiple comparisons.

An analysis of instantaneous velocities along the horizontal trajectory (reflecting the transitions between optic nerve, chiasm and proximal tract), showed some relevant differences between the crispant and non-injected embryos (Fig. 5B). For example, in the *slit2* crispants, again a greater dispersion in velocities was observed in the optic tract area, when compared to the optic nerve. In addition, both individual crispants presented some negative instantaneous velocity values accompanying sudden changes in velocity, indicative of axon misorientation events, something that was not evident in control embryos and only detected once in the double crispants (Fig. 5B). When all instant velocity data were analyzed together (Fig. 5D), a statistically significant reduction in velocity was observed in *slit2* crispants compared to non-injected control, confirming our previous observations. Interestingly, no difference to control embryos was detected for the *slit3* or the *slit2*+*slit3* crispants (Fig. 5D). In addition, an important dispersion towards negative velocities was evident particularly in *slit2* crispants (Fig. 5D).

Since the point-by-point velocity analysis shown in Fig. 5B indicated the occurrence of regional differences along the registered path, we separated the analysis of the pooled data in the two main regions, the optic nerve and the optic tract, and compared the regional distribution of velocities with those obtained for the whole path (Fig. 5D). The most remarkable observation in this regional analysis is that the main component of the observed overall instantaneous velocity reduction in *slit2* crispants is concentrated in the optic tract. In the optic nerve, there is no significant difference between the instantaneous velocities in these embryos compared to controls (Fig. 5D). In none of the cases did *slit3* or *slit2*+*slit3* crispants show a significant difference to non-injected embryos. In this experimental set up, there was no significant difference for the instantaneous velocities averaged per embryo when comparing all these situations, although the same tendency towards lower velocities was again observed in the case of *slit2* crispants, both in the whole path and in the optic tract (Fig. 5E).

### Slit2 and Slit3 differentially affect the occurrence of axon turn errors around the optic chiasm

In order to better understand the axon misorientation events mentioned above, we calculated the changes in trajectory angle with respect to the horizontal axis between timepoints for each axon (which we call “axon turns”). In the general analysis, all axon turns averaged in a value close to 0°, with a slight positive deviation of 0.5° in the control embryos, that can be explained by the anatomical curvature of the optic pathway at this level (Fig. S3). A much greater angle dispersion was observed in the *slit2* crispants, in accordance with the observed increase in misorientation events (Fig. S3). We then switched to an analysis of the absolute values of angle change, corrected by curvature, both in the whole recorded path and separating it into the optic nerve and optic tract portions (Fig. 6). When the distribution of all data points for each condition was analyzed in this way, several interesting statistically significant differences were evident, some of which we highlight here: a-turn angles were larger in *slit2* crispants than in controls and *slit3* crispants, both in the general analysis and in the optic tract; b-angles in *slit3* crispants were never larger than in non-injected embryos; c-angles significantly larger than those of controls were observed in double crispants only in the optic nerve; d-angles in *slit2*+*slit3* crispants were smaller in the optic tract than in the optic nerve, or even the whole path, and, unexpectedly, they were much smaller than those in *slit2* crispants in the optic tract (Fig. 6A).

**Figure 6.**
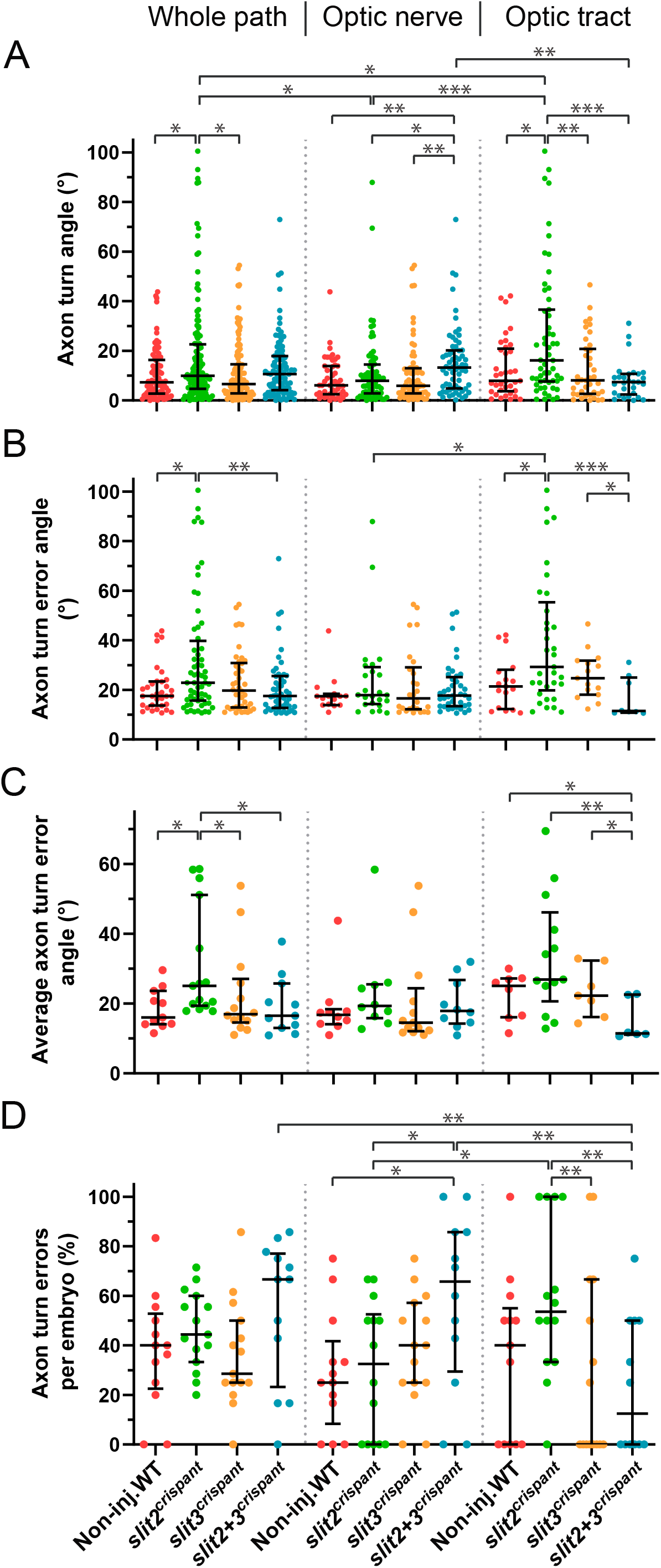
Tracking of pioneer axon growth through the optic chiasm highlights differential effects of Slit2 and Slit3 loss II: analysis of axon turn angles. The axon growth tracking data shown in Fig. 5 were analyzed to retrieve the axon turn angle for each point, relative to the previous direction, after correcting for the curvature and converting to absolute values. In all cases, the data were analyzed and compared either for the whole axon path tracked, the optic nerve portion or the optic tract portion, as noted at the top. **A**. Pooled angles for each experimental situation. **B**,**C**. Angles larger than 10.4° (“axon turn errors”) were separated and analyzed as a pool (B) or averaged per embryo (C). **D**. Number of axon turn errors per embryo, for each experimental situation, shown as a percentage of total determined angles. Median ± interquartile range; statistical significance is shown as asterisks, based on a p < 0.05 and analyzed by the Kruskal-Wallis test for multiple comparisons (only biologically relevant differences are marked).

We then proceeded to analyze the “axon turn errors”, which we defined here as points in which the turn angle was larger than the mean absolute value in control embryos (10.4°), hence excluding small misguidance events that could be frequent in all situations (Fig. 6B). In the analysis of all angle errors, either along the whole recorded path or at each region, we found very similar differences between the experimental situations as those observed in the general angle analysis shown in Fig. 6A. Interestingly, while *slit2* crispant angles were again larger, an even more marked reduction in the erroneous angle median was evident for the *slit2*-*slit3* double crispants specifically at the optic tract. These differences remained largely unchanged when averaging the erroneous angles in each embryo (Fig. 6C).

In this analysis, we were also able to determine that axons in control embryos make proportionally the same number of errors along the whole tracked path as axons in embryos deficient for *slit2* and/or *slit3* (Fig. 6D). These homogeneous results were dramatically changed, however, when analyzing each portion of the path separately. In the optic nerve, the *slit2+slit3* crispants displayed significantly more errors than controls, in accordance with a non-significant but higher median observed in the whole-path analysis (Fig. 6D). In the optic tract, on the other hand, a higher value was only evident in *slit2* crispants. Interestingly, in this region the phenotype was again significantly more severe in *slit2* single crispants than in double crispants, which presented significantly fewer errors in the optic tract than in the optic nerve or even the whole path (Fig. 6D).

## Discussion

We report a cooperative mechanism involving a simultaneous action of Slit2 and Slit3 around the optic chiasm in the zebrafish. Redundancy and complementarity of genetic pathways are expected to be widespread mechanisms in developmental processes, since they would ensure robustness (Kafri et al., 2006). An example of this is axon growth and guidance along the optic pathway, which depends on the interplay of a number of signals, including the Slit-Robo pathway. Differentiating RGCs only express Robo2 (Fricke et al., 2001), but their axons traverse a long path, in which they might encounter different, if not all, Slit secreted molecules, which in turn may also act through other receptor types. Arborization of retinal axons at the zebrafish optic tectum, for example, requires Slit1a acting both in a Robo2-dependent and -independent manner (Campbell et al., 2007). At the level of the optic chiasm, where axon crossing is finely regulated, a cooperation between Slit1 and Slit2 was described in mice (Plump et al., 2002).

Our previous observations indicated an important function of Slit2 in this region, but surprisingly, the *slit2* null mutant phenotype appeared much less severe regarding RGC axon guidance defects than the *robo2* mutant phenotype (Fricke et al., 2001; Karlstrom et al., 1996). *In situ* hybridization studies have demonstrated that two *slit* genes are expressed in the area immediately surrounding the optic chiasm in the zebrafish: *slit2* and *slit3*. Two previous reports have shown *slit2* mRNA expression very tightly surrounding the optic nerve and chiasm area (Chalasani et al., 2007; Davison and Zolessi, 2021), while *slit3* was shown by Chalasani et. al (Chalasani et al., 2007) to be highly expressed in a restricted area just caudal to the optic chiasm, with some lower expression anterior to the optic pathway (summarized in Fig. 7A). Hence, it could be possible that Slit3 is either complementing the function of Slit2 (ie: also having a function in this process in physiological conditions) or compensating for the lack of Slit2 (ie: increasing its expression in response to *slit2* gene deficiency). The quantitative mRNA expression results we presented here indicate that there are no evident changes in the expression of *slit3* in response to the null mutation of *slit2*, ruling out the compensation hypothesis. Remarkably, however, there is a small but significant increase in the expression of *slit2* in *slit3* crispant embryos. Therefore, Slit3 would not compensate for a lack of Slit2, but Slit2 could at least partially compensate for the loss of Slit3. Interestingly, this change was detected at 30 hpf, the initial stage of optic chiasm formation, and not at 48 hpf, when the process is largely completed.

**Figure 7.**
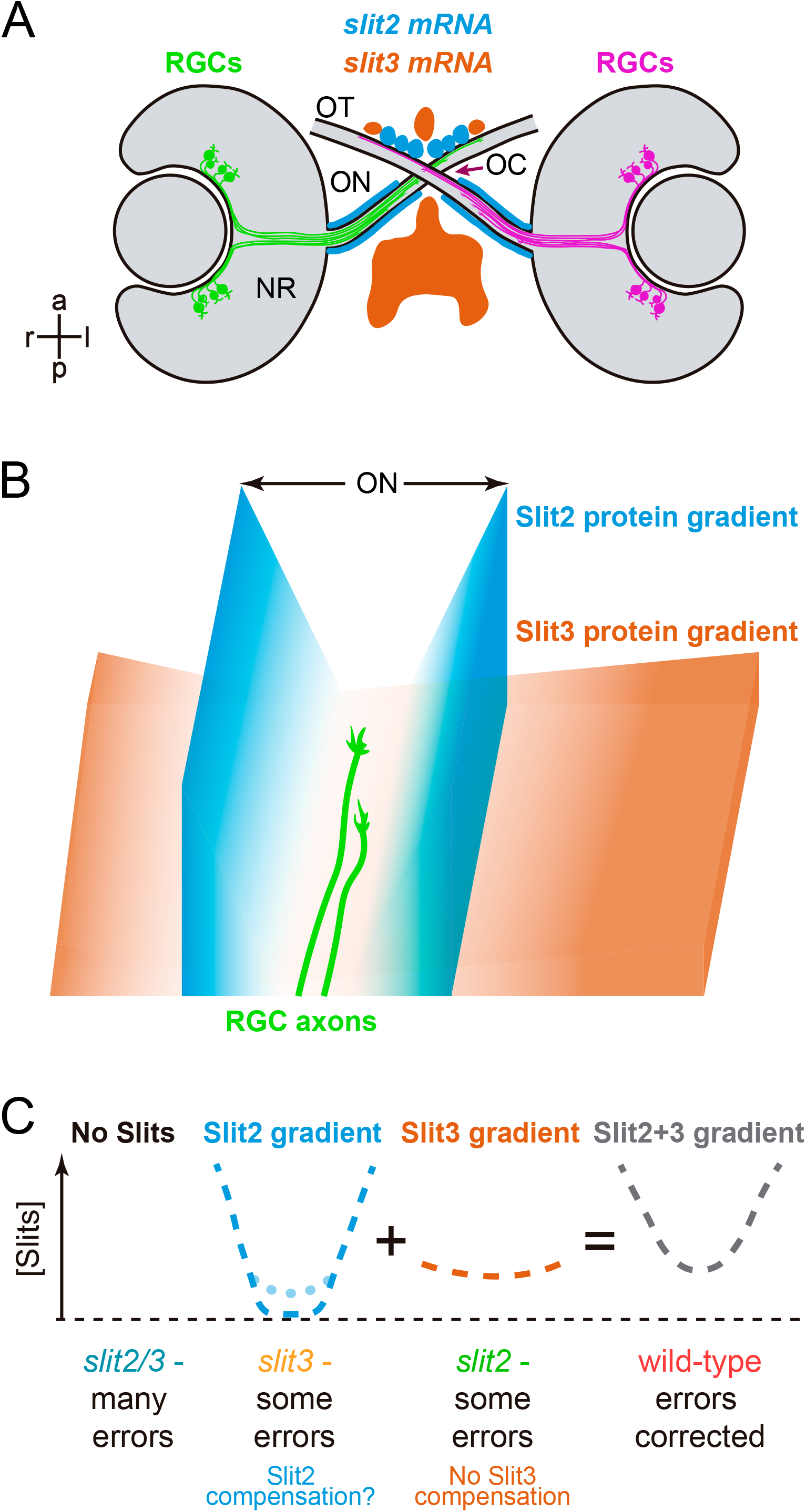
Graphical summary. **A**. Expression patterns of *slit2* and *slit3* in a wild-type zebrafish embryo, around the stage of retinal axon crossing at the chiasm, as determined by fluorescent *in situ* hybridization, and described in Chalasani et al. (Chalasani et al., 2007) and Davison and Zolessi (Davison and Zolessi, 2021). **B**. Model illustrating how Slit2 and Slit3 might be cooperating in restricting RGC axons to the optic pathway around the optic chiasm, based on the original model described by Hutson and Chien (Hutson and Chien, 2002). Slit2 expression in a narrow area tightly associated with the optic nerve and tract would generate a short but steep gradient towards the pathway center. Slit3, on the other hand, is expressed in apparent larger amounts, but at a farther distance, probably making a longer and shallower gradient around the chiasm. The summation of both gradients would result in a “U” shaped Slit, and axons would be able to grow just at the bottom of this “U”. **C**. Application of this model to the four experimental situations analyzed in the present report. If no Slits are present, retinal axons are free to grow, but make guidance errors that are not corrected, because there is no repulsive channel. Slit3 deficiency leaves only the Slit2 gradient, which is still strong enough to correct most errors; this deficiency would also slightly increase *slit2* expression, partially restoring the wild-type U gradient (dotted line). When there is a Slit2 deficiency, the remaining Slit3 gradient also allows for some uncorrected errors; there is no compensation in this case. Finally, in the wild-type situation, the full Slit (Slit2+Slit3) “U”-shaped gradient keeps axons on track by correcting all errors. NR: neural retina; OC: optic chiasm; ON: optic nerve; OT: optic tract.

A role for Slit3 in axon guidance at the zebrafish forebrain midline, both acting by itself and together with Slit2, has been previously demonstrated for other forebrain commissures. In the zebrafish supraoptic tract (SOT), for example, Slit3, and to a lesser extent Slit2, promote ipsilateral axon growth downstream of Wnt activation (Zhang et al., 2020). Moreover, knockdown of *slit2, slit3*, and *slit2*/*slit3* together resulted in defasciculation of the postoptic commissure (POC; (Barresi et al., 2005)), suggesting a channeling mechanism similar to the one we later proposed for *slit2* at the optic chiasm (Davison and Zolessi, 2021). In the research presented here, we observed that *slit3* expression disruption through the generation of mutations by the co-injection of a specific sgRNA and nCas9n, caused by itself a phenotype comparable to that of *slit2* null mutation, with the appearance of a similar proportion of cases of nerve bifurcation at the optic chiasm. In addition, *slit3* crispants displayed very minor guidance (“turn angle”) defects along the pathway, only detectable by time-lapse analysis, like was the case for *slit2* crispants. Interestingly, this increase in axon turn errors was more conspicuous in these embryos, along with a reduction in instantaneous velocity particularly in the proximal optic tract. This observation was consistent with our previous report of a severe disruption in axon segregation in this region for the *slit2* null mutant line (Davison and Zolessi, 2021).

The gross phenotype we observed in *slit2*/*slit3* double-deficient embryos included retinal axons projecting towards the ipsilateral optic tract and anteriorly into the telencephalon. Moreover, some axons appeared to follow the path of the contralateral optic nerve. Similar defects were reported for the *astray/robo2* mutant (Fricke et al., 2001; Hutson and Chien, 2002), which we reproduced in this work. Interestingly, turn angle errors made by RGC axons were equally frequent in all situations, including the wild-type embryos, while ipsilateral turns (in the direction opposite to normal growth, and evidenced here as negative instant velocity values), were present in *slit2, slit3* and *slit2+slit3* crispants, but virtually absent in wild-types. A similar phenomenon was described by Hutson and Chien (Hutson and Chien, 2002) when comparing the trajectory of RGC axons at the chiasm in wild-type and *astray/robo2* mutants by time-lapse microscopy. In addition, by following axon trajectories through lipophilic dye tracing we found ipsilateral optic tectum innervation only in *slit2+slit3* crispants, indicating that most, if not all, errors were corrected in the *slit2* or *slit3* deficient embryos. Projections to the ipsilateral optic tectum are also a hallmark of *astray/robo2* mutants (Fricke et al., 2001; Karlstrom et al., 1996). Finally, many retinal axons in *astray/robo2* mutants were observed to reach the ipsilateral tectum via the posterior commissure instead of the optic tract (Fricke et al., 2001), which could correspond to an ectopic crossing site we observed for a number of axons in *slit2+slit3* crispants, slightly posterior to the optic chiasm. Although these correlations point to the possibility of a combined action of Slit2 and Slit3 on Robo2 receptors, we cannot exclude an action on other receptors or the occurrence of more complex interactions. These could include the eventual modulation by other pathways, such as was demonstrated for SDF1-CRXR4b (Chalasani et al., 2007).

In addition to a collaborative role of Slit2 and Slit3 in assembling axon organization at the optic chiasm during the early stages of its development, our results indicate some differences in their individual functions. There are two main pieces of evidence to support this: first, there is a differential distribution of minor axon turn errors along the analyzed area (evidenced both in the spatial organization and in the dispersion of instantaneous velocity values), with errors appearing more frequently in the optic tract portion for *slit2* crispants and in the optic nerve for *slit2/3* double crispants; second, some specific phenotypic defects in the single crispants appeared to be corrected in the double deficient embryos, like the distal turn errors in *slit2* crispants. The different expression patterns for these two genes could at least partly explain these differences (see Fig. 7A). Since retinal axons are known to respond differently to guidance cues depending on their position, it would be interesting to determine whether the observed differential defects are related to actions on specific axon populations. A role like this was proposed for HSPGs in regulating dorso-ventral sorting in the zebrafish optic tract through selective axon degeneration (Poulain and Chien, 2013). Similarly, Slit molecules could be regulating naso-temporal axon segregation at the optic chiasm and proximal tract, like we described for *slit2* morphants (Davison and Zolessi, 2021), albeit most probably through error correction.

Based on these previous data and our present observations, we suggest a hypothetical model by which Slit2 would have a “channeling” role, as proposed by Chi-Bin Chien and colleagues several years ago (Hutson and Chien, 2002; Rasband et al., 2003), with steep gradients surrounding the path of optic axons before, across and just after the optic chiasm. Slit3, on the other hand, would generate a smoother and wider gradient, eventually preventing the axons from entering into the brain as they cross at the optic chiasm, and nevertheless collaborating with Slit2 in preventing axon turn errors at the midline (see a diagram in Fig. 7B). Together, they would generate a “U” shaped Slit gradient to keep retinal axons on track. Loss of Slit2 would only leave the shallow Slit3 gradient, leading to relatively important defects in axon organization particularly in the proximal optic tract; loss of Slit3 would, in turn, leave the marked Slit2 gradient intact, with the possibility of an increased local expression due to genetic compensation, and leading to milder defects overall; the simultaneous loss of Slit2 and Slit3 would lead to a Robo2-like phenotype, due to a complete absence of Slits in the area surrounding the optic chiasm (schematic representation in Fig. 7C).

It is interesting to note that Slit2 and Slit3 in the zebrafish could be acting in ways comparable to those described for Slit1 and Slit2, respectively, in mice (Plump et al., 2002), as in this species Slit1 is expressed in a pattern tightly surrounding the optic pathway proximal to the chiasm, and Slit2 is expressed in a larger area located in the anterior midline, at some distance (Erskine et al., 2000). In mice, like in most mammalian species, there is some degree of binocular vision, and some retinal axons do not cross at the optic chiasm, but project ipsilaterally. It is tempting to speculate that the use of different Slit factors, with differential expression patterns, all acting on Robo2, could be part of an evolutionary mechanism for adaptation to different degrees of binocular or non-binocular vision, which appears more related to the particular vertebrate species habits than to phylogenetic constraints.

## Acknowledgements

The authors thank William A. Harris, University of Cambridge, for support with lab space and materials, as well as fruitful discussion; Jon Clarke, King’s College London, for sending the *astray* mutant line; Kristen Kwan, University of Utah, for providing Tol2-kit plasmids; Casandra Carrillo and Gisell González, Zebrafish Lab, Institut Pasteur Montevideo, for fish maintenance and care. The authors gratefully acknowledge the Advanced Bioimaging Unit at the Institut Pasteur Montevideo for their support and assistance in the present work.

## Author contributions

CD designed and performed the experimental manipulations, analyzed the data and wrote the manuscript; GB designed, performed and analyzed the quantitative RT-PCR experiments, and helped writing the manuscript; FRZ directed the research, designed part of the experiments, analyzed part of the data and wrote the manuscript.

## Data availability statement

The data that support the findings of this study, as well as original materials, are available from the corresponding author upon reasonable request.

## Figure legends

**Figure S1.**
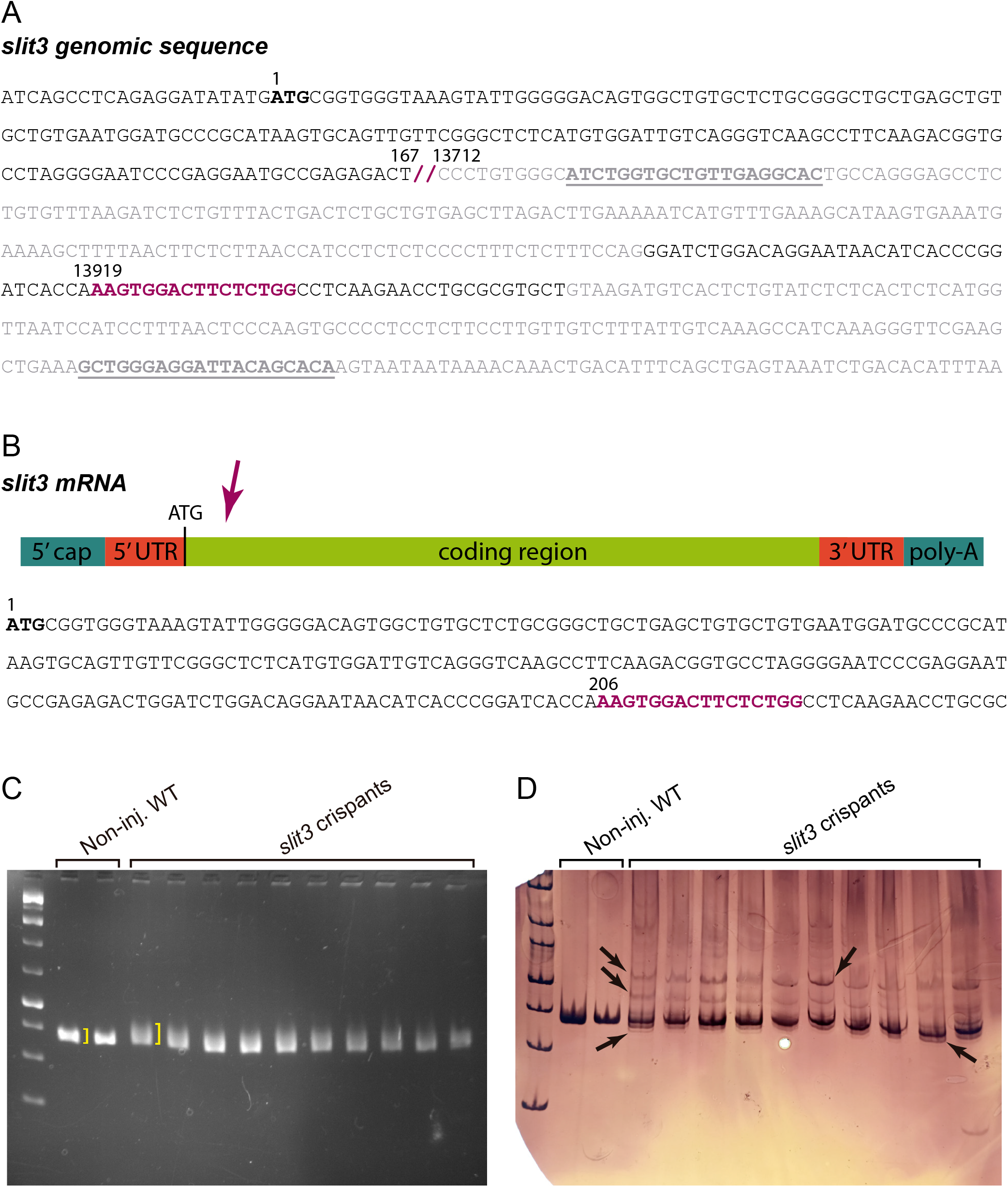
Injection of sgRNA 206 together with nCas9n induces indels in the *slit3* gene target site. **A**. Partial sequence corresponding to the zebrafish *slit3* gene. Exons are represented in black, while introns are coloured in grey. The initiation codon (ATG) is marked in bold. The entirety of the first exon is shown, while only the 3’-most region of the first intron can be observed, with the double bar indicating an unshown sequence belonging to the first intron. The *slit3* 206 sgRNA-binding site is shown in bold and coloured in purple, while the sequences complementary to the primers used for genotyping are underlined. **B**. Schematic representation of the *slit3* mRNA. The arrow indicates the approximate site of expected mutagenesis. The coding sequence of *slit3* is also shown, with the sgRNA binding site indicated in bold purple. **C**,**D**. Agarose (**C**) and polyacrylamide (**D**) gel electrophoresis analyses showing PCR products obtained from genomic DNA. The same samples were run in both gels, with each lane corresponding to an individual embryo. Two non-injected wild-types and ten *slit3* crispants were analyzed. Brackets in the agarose gel indicate the difference in band width between wild-type and crispant embryos; arrows in the polyacrylamide gel indicate extra bands, not observed in wild-types.

**Figure S2.**
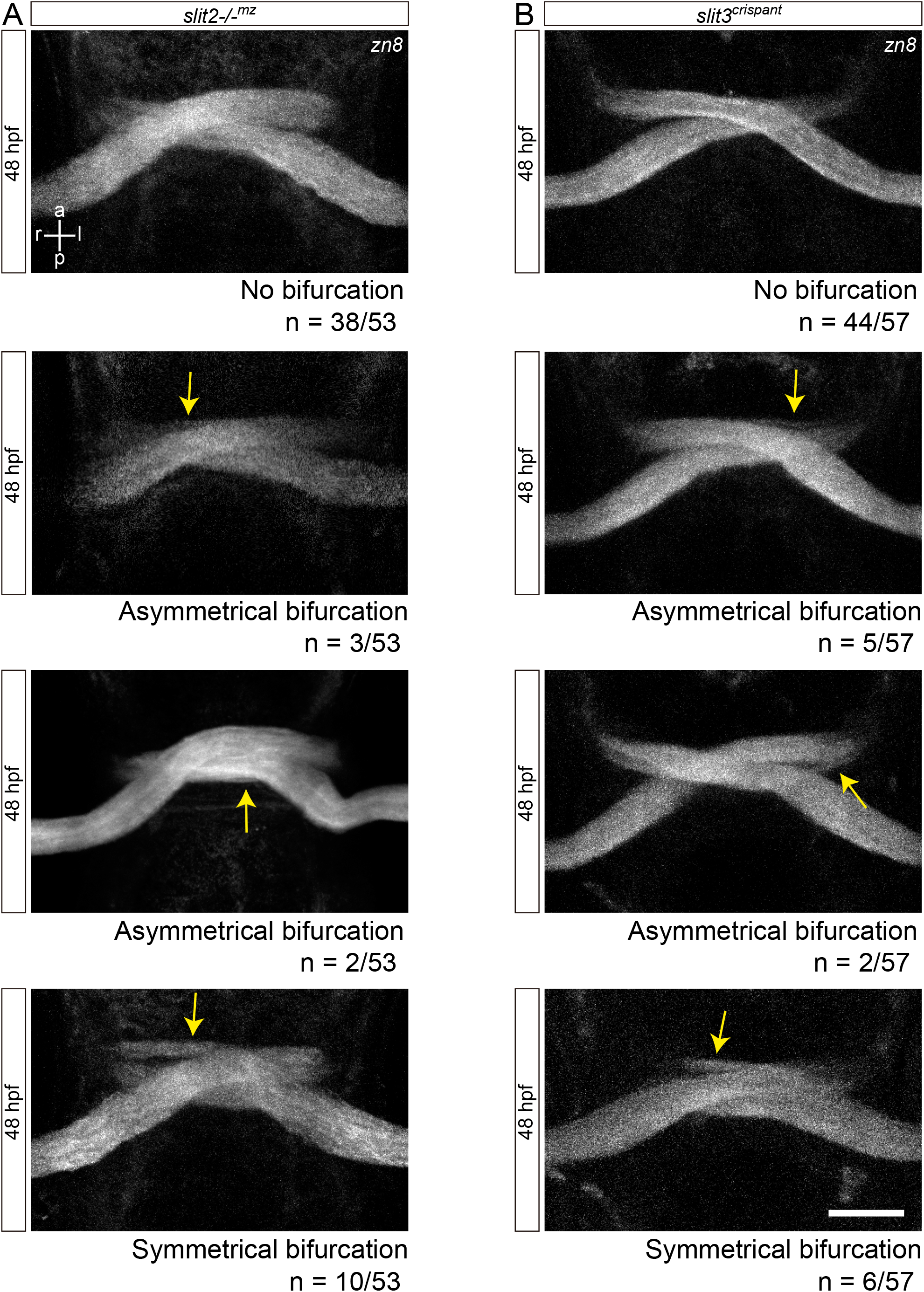
Analysis of the bifurcation phenotype in *slit2-/-*^*mz*^ and *slit3* crispant embryos. **A**,**B**. Maximum intensity z-projections of the optic chiasm region of 48 hpf embryos immunostained to label RGCs (zn8 antibody; ventral view). Four different phenotypes can be seen in *slit2-/-*^*mz*^ (**A**) and *slit3* crispant (**B**) embryos: no bifurcation, asymmetrical bifurcation with a thin anterior branch, asymmetrical bifurcation with a thin posterior branch, and symmetrical bifurcation with branches of similar thickness. The number of cases observed for each phenotype is shown. Arrows indicate the site of bifurcation. Scale bar: 30 μm.

**Figure S3.**
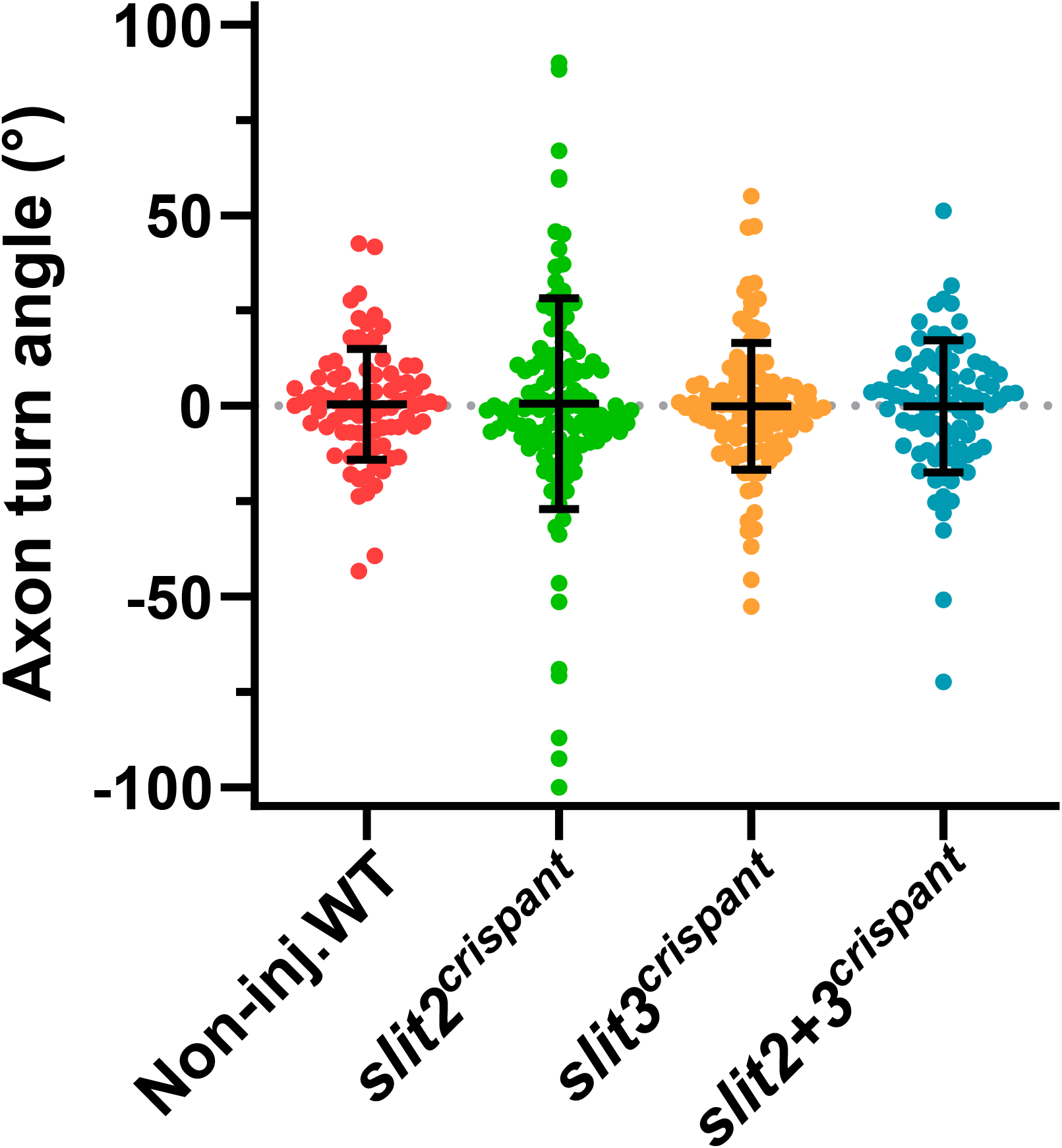
Axon turn angles across the optic chiasm. The angle with respect to the horizontal axis (medio-lateral) was determined for each time point, from each axon on the tracking analysis shown in Fig. 5A. All angles obtained for each experimental condition were plotted together in this graph, to compare general angle distributions and mean ± SD. Positive angles indicate dorsal turn, and negative angles ventral turn. Angle means are not significantly different (Brown-Forsythe and Welch test), and the mean value for the non-injected wild-type embryos is around 0.5°, following the curvature of the ventral diencephalon.

**Video S1. Simultaneous loss of Slit2 and Slit3 causes severe axon guidance defects at the optic chiasm**. Z-stacks showing the optic chiasm of non-injected wild-type, *slit2-/-* ^*mz*^, *slit3* crispant and *slit2-/-*^*mz*^;*slit3* crispant embryos at 48 hpf, after immunostaining with zn8 antibody. In non-injected wild-type embryos, one optic nerve can be seen crossing anteriorly to the contralateral nerve, while in *slit2-/-*^*mz*^ and *slit3* crispant embryos, one of the optic nerves splits and surrounds the contralateral nerve in ≈25% of the cases. *slit2-/-*^*mz*^;*slit3* crispant embryos display more severe defects, which include projections to the anterior telencephalon. The stack sequence is shown from the ventral to the dorsal region of the embryo. Corresponding to Fig. 1A.

**Video S2. Simultaneous loss of Slit2 and Slit3 causes retinal axon misprojections at the optic chiasm**. 3D projections of z-stacks of the optic chiasm of non-injected wild-type, *slit2-/-*^*mz*^, *slit3* crispant and *slit2-/-*^*mz*^;*slit3* crispant embryos observed at 48 hpf, where RGC axons from both eyes were labeled anterogradely with either DiI or DiO. In non-injected wild-type embryos, one of the optic nerves can be seen crossing anteriorly and slightly ventral to the contralateral nerve, while remaining physically separate. In *slit2-/-*^*mz*^ and in *slit3* crispant embryos, one of the optic nerves can be seen splitting into two groups of axons which surround the contralateral nerve. More severe defects can be seen in *slit2-/-*^*mz*^;*slit3* crispant embryos, including projections to the ipsilateral optic tract and the contralateral optic nerve. Corresponding to Fig. 2.

**Video S3. RGCs in *slit2/slit3* double crispants partially project to the ipsilateral optic tectum**. 3D projection of z-stack of the optic tectum of an *atoh7*:GFP transgenic *slit2+slit3* crispant 5 dpf larva, where RGC axons from one eye were labeled anterogradely with DiI. The tectum ipsilateral to the labeled eye is shown, where DiI-labeled axons can be seen. Corresponding to Fig. 3D,d’.

**Video S4. Retinal axons from *slit2, slit3* and *slit2/slit3* double crispants present navigation errors around the midline**. Confocal time-lapse (4D) maximum intensity projection images of the optic chiasm region from *atoh7*:GFP transgenic embryos, acquired every 15 min from 30 hpf. A representative non-injected wild-type, *slit2* crispant, *slit3* crispant and *slit2+slit3* crispant are shown. At the end, a full stack reconstruction of the methyl green-counter-stained embryos 16.5 h after the beginning of the time-lapse experiments is shown. Ipsilateral turns and minor pathfinding errors can be seen in *slit2* crispant, *slit3* crispant and *slit2+slit3* crispants, but not non-injected wild-types. At the end of the experiment, these optic chiasms show clear defects in the sorting of axons coming from each eye. Corresponding to Fig. 4, see also Videos S5-8.

**Video S5. Retinal axons from *slit2, slit3* and *slit2/slit3* double crispants present navigation errors around the midline II**. Confocal time-lapse (4D) maximum intensity projection images of the optic chiasm region from an *atoh7*:GFP transgenic embryo injected with *slit2* sgRNA/nCas9n, acquired every 15 min from 30 hpf. At the end, a full stack reconstruction of the methyl green-counter-stained embryo 16.5 h after the beginning of the time-lapse experiments is shown. Corresponding to Fig. 4, see also Videos S4,6-8.

**Video S6. Retinal axons from *slit2, slit3* and *slit2/slit3* double crispants present navigation errors around the midline III**. Confocal time-lapse (4D) maximum intensity projection images of the optic chiasm region from an *atoh7*:GFP transgenic embryo injected with *slit3* sgRNA/nCas9n, acquired every 15 min from 30 hpf. At the end, a full stack reconstruction of the methyl green-counter-stained embryo 16.5 h after the beginning of the time-lapse experiments is shown. Corresponding to Fig. 4, see also Videos S4,5,7,8.

**Video S7. Retinal axons from *slit2, slit3* and *slit2/slit3* double crispants present navigation errors around the midline IV**. Confocal time-lapse (4D) maximum intensity projection images of the optic chiasm region from an *atoh7*:GFP transgenic embryo injected with *slit2+slit3* sgRNA/nCas9n, acquired every 15 min from 30 hpf. At the end, a full stack reconstruction of the methyl green-counter-stained embryo 16.5 h after the beginning of the time-lapse experiments is shown. Corresponding to Fig. 4, see also Videos S4-6,8.

**Video S8. Retinal axons from *slit2, slit3* and *slit2/slit3* double crispants present navigation errors around the midline V**. Confocal time-lapse (4D) maximum intensity projection images of the optic chiasm region from an *atoh7*:GFP transgenic embryo injected with *slit2+slit3* sgRNA/nCas9n, acquired every 15 min from 30 hpf. At the end, a full stack reconstruction of the methyl green-counter-stained embryo 16.5 h after the beginning of the time-lapse experiments is shown. Corresponding to Fig. 4, see also Videos S4-7.

## SUPPLEMENTARY TABLES

**Table 1.**
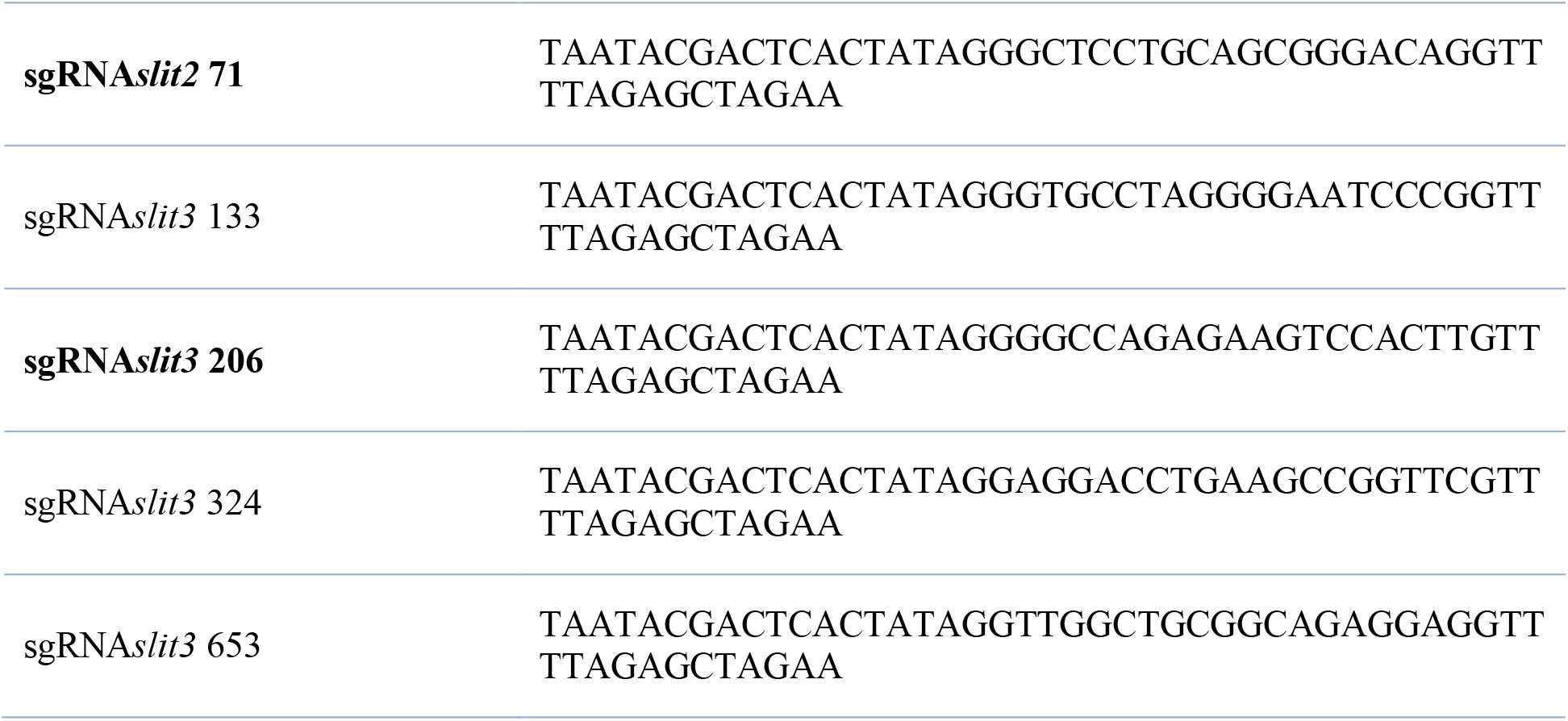
Single-guide RNA sequences used for CRISPR/Cas9 mutation.

**Table 2.**
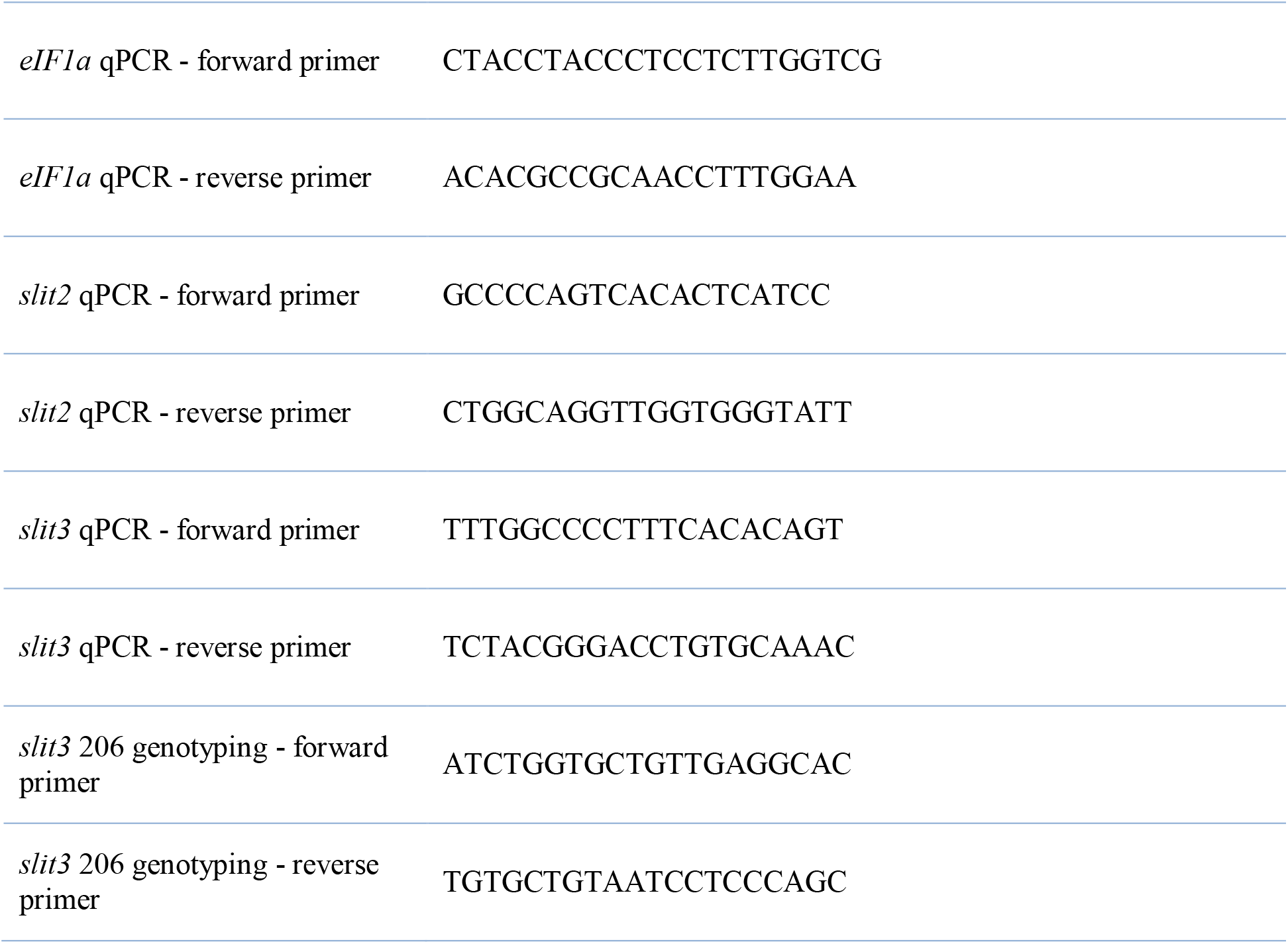
Primer sequences used for quantitative PCR analyses.

